# Recovery of cryo-injured rabbit urethras by biofabricated C-shaped adipose-derived mesenchymal cell structures

**DOI:** 10.1101/2020.04.02.022715

**Authors:** Sudha Silwal Gautam, Tetsuya Imamura, Mitsuru Shimamura, Tomonori Minagawa, Masaki Nakazawa, Teruyuki Ogawa, Osamu Ishizuka

**Affiliations:** Department of Urology, Shinshu University School of Medicine, Nagano, Japan; Cyfuse Biomedical K.K., Tokyo, Japan

**Keywords:** Adipose-derived stem cell, Biofabrication, Cryo-injury, Urethra, Skeletal muscle, Smooth muscle

## Abstract

Urethral tissue damage can cause stress urinary incontinence or urinary retention, and there are few long-term, effective treatments. For structural and functional recovery of urethral tissues, we designed and constructed C-shaped structures composed of adipose-derived mesenchymal cells (AMCs) by using a three-dimensional (3D)-bioprinter system, which could be transplanted in damaged urethras without obstructing the lumen. We determined if transplantation of the biofabricated structures could reconstruct the urethral tissues. AMCs were harvested from rabbits, cultured and labeled with PKH26 to form spheroids. The spheroids were assembled on a custom-designed C-shaped support by a 3D bioprinter. Urethras of rabbits were cryo-injured by spraying with liquid nitrogen for 20 seconds, incised and biofabricated structure was autologously transplanted. Control rabbits were treated similarly but without transplantation structure. Two and four weeks after surgery, the control urethras were partially constricted; however, the structure-transplanted urethras were patent. The AMCs within the structures differentiated into skeletal muscle, smooth muscle, nerve, or endothelial cells. Some cells contained growth factors and cytokines. Therefore, biofabricated C-shaped AMC structures have potential to be an effective treatment for urethral recovery.

**Summary Statement:** Three-dimensional bioprinter was used to biofabricate novel C-shaped structures composed of adipose-derived mesenchymal cells (AMCs) and have the potential to be an effective treatment for urethral recovery.

## Introduction

Patients with urethral tissue damage due to aging often complain about stress urinary incontinence (SUI), which is one manifestation of urine storage dysfunction. Also, urethral injury or post-operation stricture can induce urinary retention, which is a voiding dysfunction. Long-term, safe, and effective treatments for the urethral tissue damage have not been developed. Recently, mesenchymal stem cell-based therapies have been attempted as treatments for SUI. In our previous study, we showed that injection of autologous adipose- (Silwal Gautam et al., 2014) or bone marrow-derived mesenchymal cells (Imamura et al., 2011) into the cryo-injured urethra of rabbits reconstructed the functional urethra. Clinical research suggests that injection of autologous adipose-derived cells into the urethra of male patients with SUI has the potential to provide significant benefits (Aragon et al., 2018; Yamamoto et al., 2012). Thus, autologous cells including somatic mesenchymal stem cells, such as adipose- or bone marrow-derived mesenchymal stem cells have the potential to repair injury-damaged urethras and thus warrant further study.

However, there are some limitations to the cell injection methods. The cells can be damaged during the harvesting and handling procedures. The most serious limitation is the low survival rate of the injected cells in the recipient tissues. Previously, we reported that biofabricated structures improve cell survival rates during harvesting and handling and in the recipient tissues (Imamura et al., 2018; Moldovan et al., 2017). In this study, we designed novel biofabricated structures composed of adipose-derived mesenchymal cells (AMCs) and constructed them with a three-dimensional (3D)-bioprinter. After construction, the structures were inserted into the injured urethras of rabbits where they promoted healing of the damaged tissues. By design, the structures were C-shaped to avoid blocking the lumen of the urethra and potentially impeding the flow of urine. Here we report the biofabrication methodology, the transplantation of the C-shaped structures into the damaged urethras, and the reconstruction of the damaged urethral tissues through cellular proliferation and differentiation. We also report evidence of paracrine stimulation through the presence of growth factors and cytokines within transplanted structure and adjacent tissues.

## Results

### Characterization of AMCs within biofabricated C-shaped structures

Before urethral transplantation, the PKH26-labeled AMCs within the biofabricated structures (Fig. 2A) were positive for mesenchymal cell marker STRO-1 antibody (Fig. 2B, C). They also expressed integrin β1 (Fig. 2D-F) and N-cadherin (Fig. 2G-I) in the structures. However, other markers, such as those for skeletal muscle (myoglobin), smooth muscle (smooth muscle actin), nerve (S100), and vascular endothelial cell (vWF), were not present.

**FIGURE 1.**
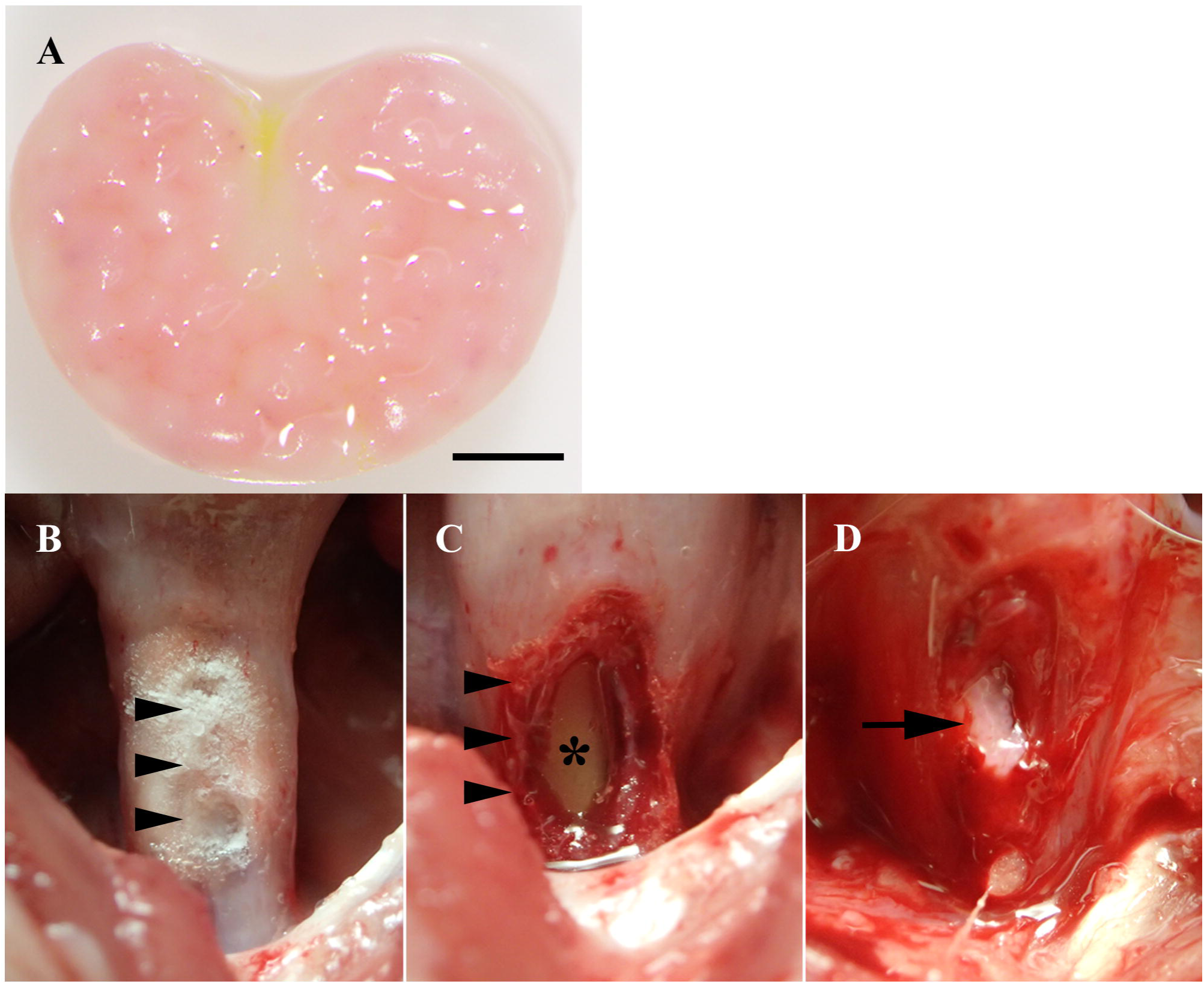
Autologous C-shaped AMC structure, urethral cryo-injury, and transplantation into the injured urethra. **(A)** Biofabricated C-shaped AMC structure after removal from microneedle array. The structure was approximately 3.5 (wide)×3.5 (depth)×1.0 (high) mm, Scale bar: 1 mm. **(B)** The intact urethra was sprayed with liquid nitrogen for 20 second (arrowheads). **(C)** A small incision of approximentally 5 mm was created around the cyroinjured regions (arowheads). Asterisk, catheter in the urethral lumen. **(D)** Biofabriacted C-shaped AMC structure (arrow) was implanted into the incised urethra.

**FIGURE 2.**
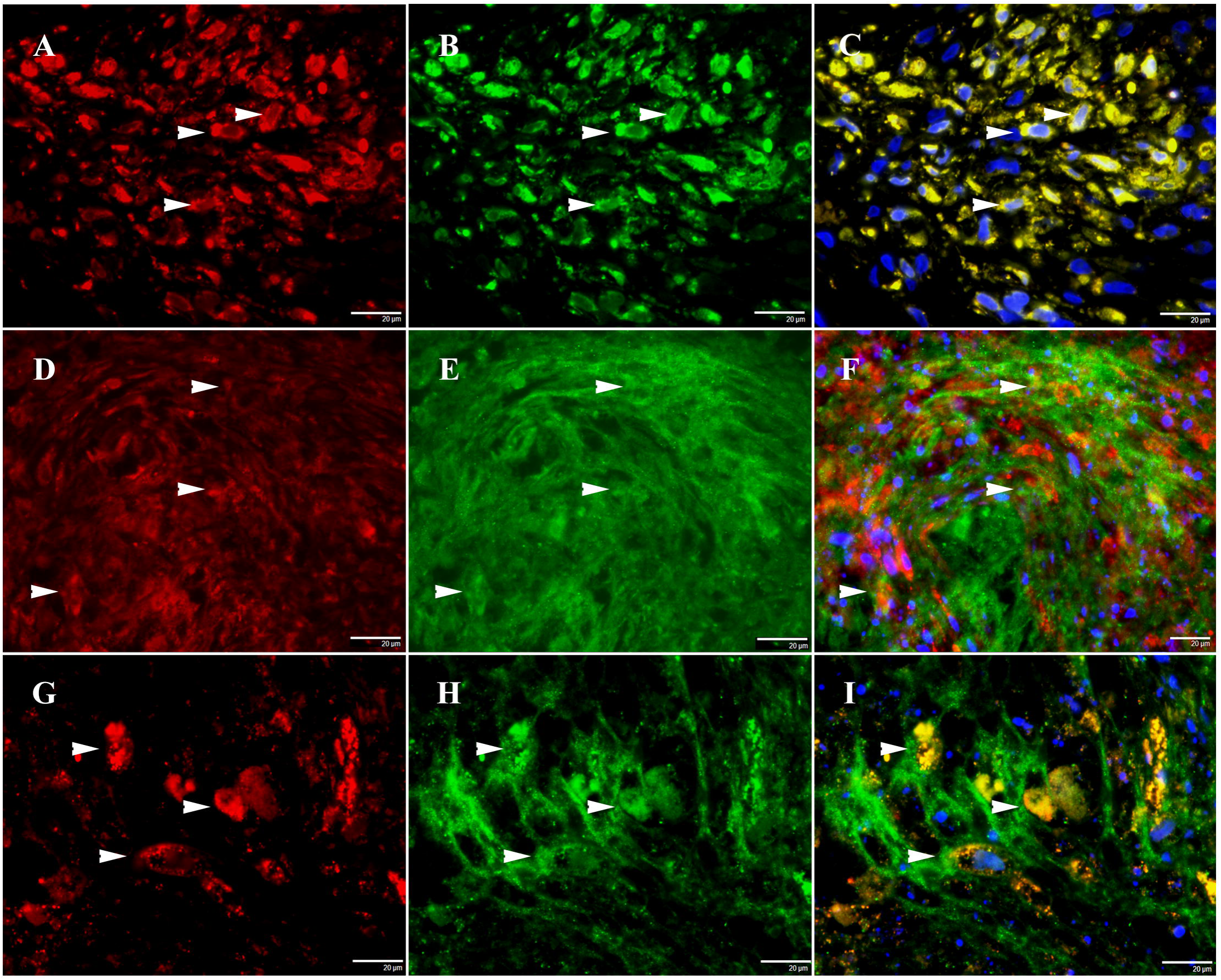
Expression of mesenchymal and extracellular matrix markers of AMCs within biofabricated structures. **(A-C)** The PKH26-labeled AMCs within the biofabricated structures (A, red, arrowheads) were positive for the mesenchymal cell marker STRO-1 (B, green, arrowheads; C, merged image, arrowheads). **(D-F)** The PKH26-labeled AMCs (D, red, arrowheads) were positive for the extracellular marker integrin (E, green, arrowheads; F, merged image, arrowheads). **(G-I)** The PKH26-labeled AMCs (G, red, arrowheads) were positive for the extracellular marker cadherin (H, green, arrowheads; I, merged image, arrowheads). Blue: DAPI-labeled nuclei, Scale bar: 20 μm.

### Recovery of urethral tissues after transplantation of biofabricated C-shaped structures

Two weeks after the sham operation, the cryo-injured and incised regions of the sham control urethras had developed strictures (Fig. 3A). Furthermore, in the sham-operated regions, there were skeletal and/or smooth muscle cells within the urethral tissues, but they were sparsely distributed and disorganized (Fig. 3B). In contrast, at two weeks after surgery, the urethras transplanted with a C-shaped biofabricated structure were wider and straight compared to the controls (Fig. 3C). The urethral tissues in the biofabricated structure-transplanted regions also had skeletal and smooth muscle cells (Fig. 3D).

**FIGURE 3.**
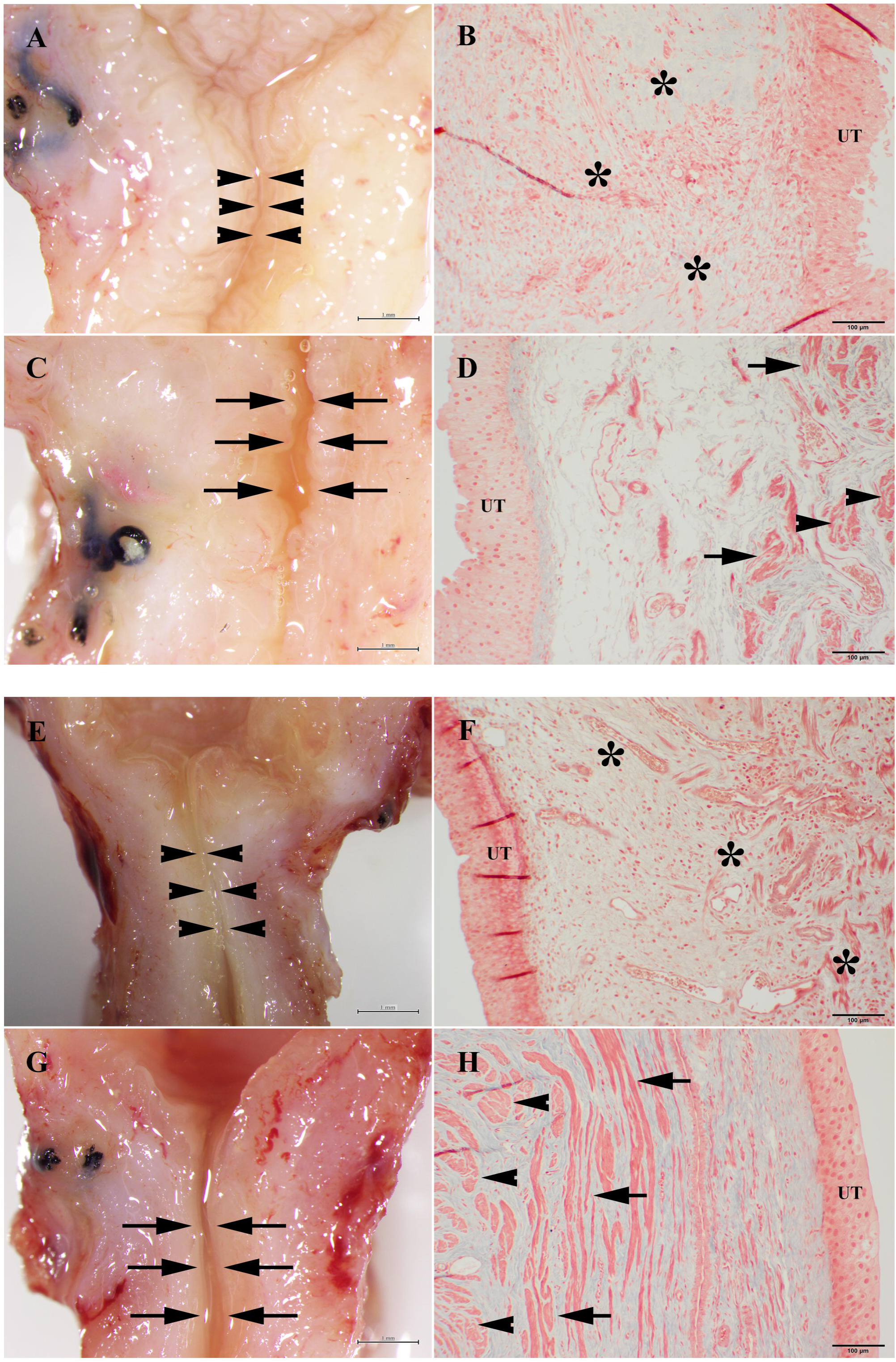
Urethral tissues transplanted with and without biofabricated structures at two and four weeks after surgery. **(A)** At two weeks after the sham operation, the gross appearance of the control urethra was narrowed and somewhat tortuous (arrowheads). **(B)** The skeletal muscle cells and/or smooth muscles cells within the control urethra were sparsely distributed and disorganized (asterisks). Scale bar: 100 μm. **(C)** At two weeks after transplantation of a AMC biofabricated structure, the gross appearance of the urethra was wider and straighter compared to the sham-operated control (arrows). **(D)** At two weeks, a urethra transplanted with a biofabricated structure had skeletal (arrows) and smooth muscle (arrowheads) cells. **(E)** At four weeks after sham surgery, the urethra had strictures that were similar to those at two weeks after surgery (arrowheads). **(F)** The walls of the sham-operated controls had some skeletal and smooth muscle cells, but they did not form typical muscle layers (asterisk). **(G)** At four weeks after surgery, the structure-transplanted urethra had a wider lumen (arrows) that was straighter than in the controls. **(H)** At four weeks after structure transplantation, the uretha had numerous skeletal (arrows) and smooth muscle (arrowheads) cells that were organized into layered tissue structures. Scale bar: gross images (A, C, E, and G): 1 mm; Masson trichrome staining (B, D, F, and H): 100 μm; UT, urothelium.

At four weeks after the sham operation, the control rabbits continued to have urethral strictures that were similar to those seen at two weeks after surgery (Fig. 3E). While the controls had some skeletal and smooth muscle cells, they were not organized into muscle layers (Fig. 3F). However, the biofabricated structure-transplanted urethras, at four weeks after surgery, did not have urethral strictures or other obstructions (Fig. 3G). Similar to the findings at two weeks, there were numerous skeletal and smooth muscle cells, but these were now organized into layered muscle-like structures (Fig. 3H).

### Survival and differentiation of AMCs within the biofabricated structures after transplantation

At both two and four weeks after transplantation, PKH26, the red fluorescent dye embedded in the membranes of the AMCs before spheroid formation, was present in AMCs of the transplanted structures. At four weeks after transplantation, some of the PKH26-labeled cells were positive for antibodies to the skeletal muscle marker myoglobin (Fig. 4A-C) or the smooth muscle marker SMA (Fig. 4D-F). Also, some PKH26-labeled cells were positive for antibodies to the immature skeletal muscle cell marker desmin (Fig. 4G-I) or the myoblast progenitor cell marker pax7 (Fig. 4J-L). Moreover, some of the PKH26-labeled cells migrated from the transplanted structures into the surrounding urethral regions (data not shown).

**FIGURE 4.**
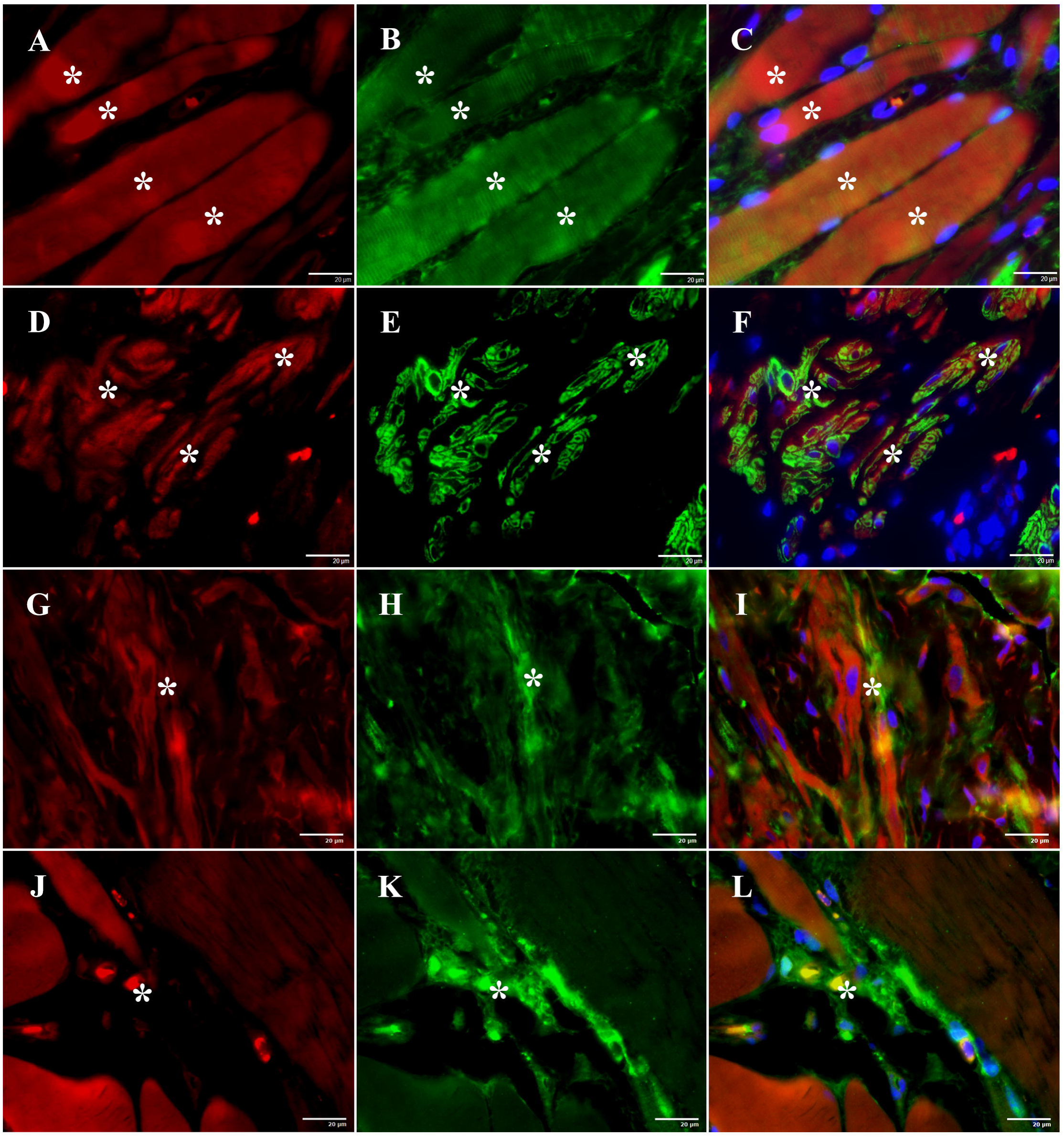
Differentation of AMCs within the transplanted biofabricated structures into muscle marker-positive cells at four weeks after transplantation. **(A-C)** The PKH26-labeled cells (A, red, asterisks) were positive for the skeletal muscle marker myoglobin (B, green, arrowheads; C, merged image, asterisks). **(D-F)** The PKH26-labeled cells (D, red, asterisks) were positive for the smooth muscle marker SMA (E, green, asterisks; F, merged image, asterisks). **(G-I)** The PKH26-labeled cells (G, red, asterisks) were positive for the immature skeletal muscle cell marker desmin (H, green, asterisks; I, merged image, asterisks), **(J-L)** The PKH26-labeled cells (J, red, asterisk) were positive for the myoblast progenitor cell marker pax7, (K, green, asterisk; L, merged image, asterisks). Blue: DAPI-labeled nuclei, Scale bar: 20 μm.

Some of PKH26-labeled AMCs were positive for the nerve cell marker S100 (Fig. 5A-C) or the afferent nerve marker CGRP (Fig. 5D-F). In addition, other PKH26-labeled AMCs were positive for the vascular endothelial cell marker vWF (Fig. 5G-I) at both two and four weeks after transplantation. The vWF positive cells with PKH26 were integrated within the blood vessel-like structures within the transplanted biofabricated structure (Fig. 5I).

**FIGURE 5.**
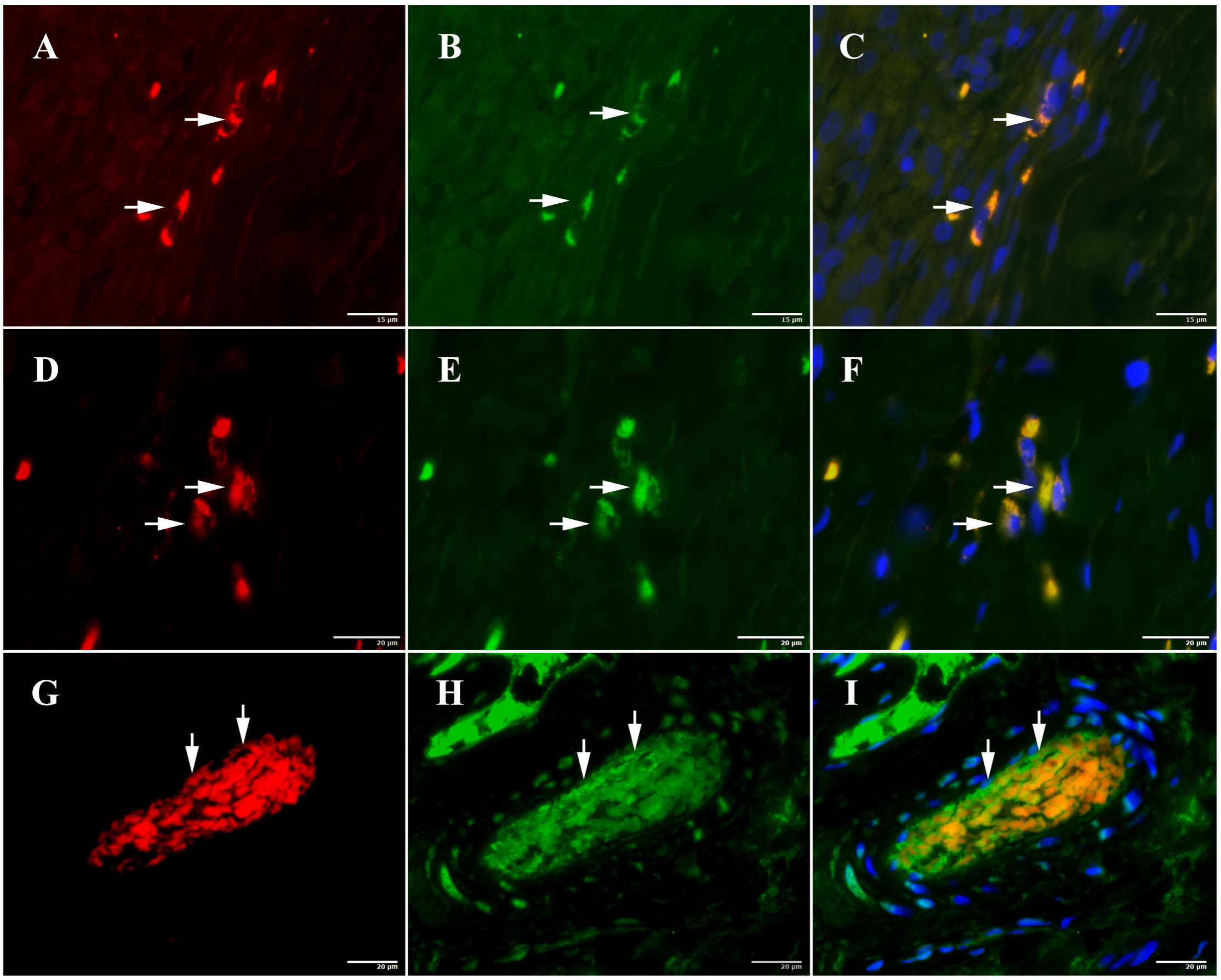
Differentation of AMCs within transplanted biofabricated structures into nerve or vascular endothelial marker-positive cells at four weeks after transplantation. **(A-C)** The PKH26 labeled cells (A, red arrows) were positive for the nerve cell marker S100 (B, green, arrows; C, merged image, arrows). Scale bar: 15 μm. **(D-F)** The PKH26 labeled cells (D, red arrows) were positive for the afferent nerve marker CGRP (E, green, arrows; F, merged image, arrows). Scale bar: 20 μm. **(G-I)** The PKH26 labeled cells (G, red arrows) were positive for the vascular endothelial marker vWF (H, green, arrows; I, merged image, arrows). Blue: DAPI-labeled nuclei, Scale bar: 20 μm.

### AMCs growth factors and cytokines within the biofabricated structures after transplantation

At both two and four weeks after the sham operation, some cells in the control regions contained VEGF (Fig. 6A). Similarly, some cells in the transplantation regions were also positive for VEGF (Fig. 6B). In addition, the cells labeled with PKH26, indicating that they were derived from the AMCs, also contained VEGF (Fig. 6B). Another growth factor, NGF, was also detected within some cells in the both control (Fig. 6C) and transplantation (Fig. 6D) regions. The control and transplanted regions also contained the cytokines TGF-β1 (Fig. 6E, F) and TNF-α (Fig. 6G, H). In the transplanted regions, some PKH26-labeled AMCs were also positive for NGF (Fig. 6D), TGF-β1 (Fig. 6F), and TNF-α (Fig. 6H).

**FIGURE 6.**
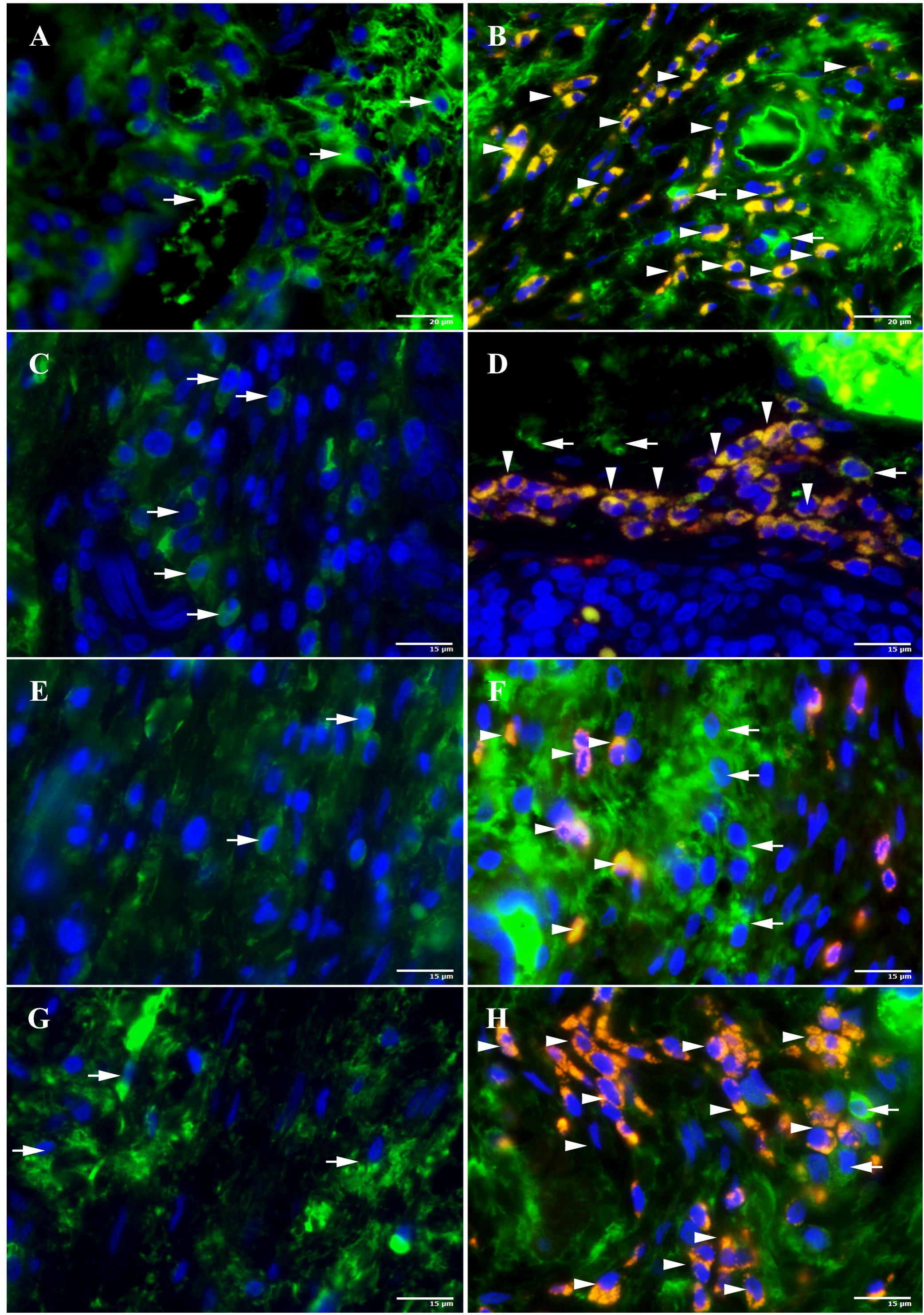
Growth factors and cytokines within the sham-operated control and transplanted regions and in the PKH26-labeled AMCs at four weeks after surgery. The sham-operated control and transplanted regions were identified macroscopically by the presence of suture marks. **(A, B)** Within the control (A) and transplanted regions (B), some cells contained VEGF (green, arrows). In addition, the PKH26-labeled AMCs also contained VEGF (B, yellow, arrowheads). **(C-H)** Similarly, some cells within the control (C, E, G) and transplanted regions (D, F, H) contained NGF (C and D, green, arrows), TGF-β1 (E and F, green, arrows), and TNF-α (G and H, green, arrows). Some of the PKH26-labeled AMCs in the transplanted regions contained NGF (D, yellow, arrowheads), TGF-β1 (F, yellow, arrowheads) and TNF-α (H, yellow, arrowheads). Blue: DAPI-labeled nuclei, Scale bar: A, B: 20 μm; C – H: 15 μm.

### Histological changes in biofabricated structure-transplanted urethral tissues

We observed cryo-injured urethral tissues at the borders of the wound in sham-operated regions and between the biofabricated AMC structures and the recipient tissues in the transplanted regions. At two and four weeks after sham surgery in which no biofabricated structure was transplanted, the urethral wound site contained collagen fibers in the extracellular matrix along with damaged skeletal muscle and smooth muscle layers (Fig. 7A). In contrast, in urethras containing the transplanted biofabricated structures, the collagen fibers were located between the recovering skeletal and smooth muscle layers (Fig. 7B). The control urethras had numerous fibroblast P4HB-positive cells, an indicator of hypoxia, within the collagen fibers (Fig. 7C) compared to the structure-transplanted urethras (Fig. 7D). The control urethras also had numerous HIF1α-positive cells, another indicator of hypoxia (Fig. 7E), and apoptotic cells (Fig. 7G). In contrast, the structure-transplanted urethra had few HIF1α-positive or apoptotic cells (Fig. 7F, H).

**FIGURE 7.**
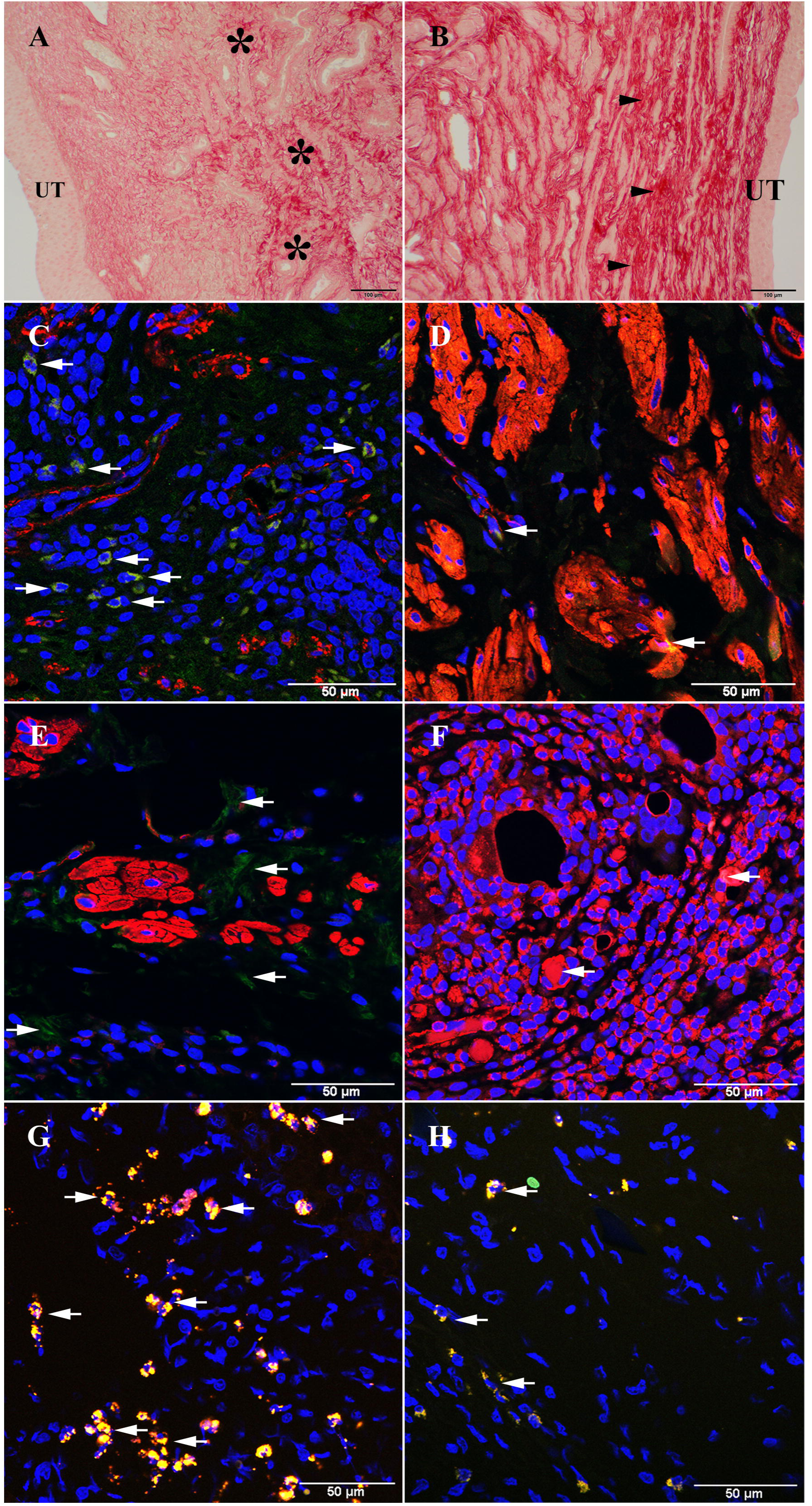
Histological changes in sham-operated control urethral tissues and biofabricated structure-transplanted tissues at four weeks after surgery. **(A-B)** Within the control regions (A), the collagen fibers had invaded the spaces of the damaged skeletal muscle or smooth muscle layers (asterisks). In contrast, in the structure-transplanted regions (B), the collagen fibers were located between the recovering skeletal and smooth muscle layers (arrowheads). Picrosirus red staining for collagen. Scale bar: 100 μm. **(C-D)** In the control urethra, some cells within the collagen fibres were positive for P4HB antibody, an indicator of hypoxia (C, green, arrows). However, the transplanted regions had few P4HB-positive cells (D, green or yellow, arrows). Red: SMA-positive smooth muscle cells. **(E-F)** The control regions had numerous HIF1α-positive cells, another indicator of hypoxia (E, green, arrows). In contrast, the transplanted regions had very few HIF1α-positive cells. (F, yellow, arrows). Red: SMA-positive smooth muscle cells. **(G-H)** The control regions had numerous apoptotic cells (G, green, caspase-independent apoptotic cells; red, caspase-dependent apoptotic cells; yellow, overlapping red-labeled caspase-dependent and green-labeled caspase-independent apoptotic cells, arrows). However, in the transplanted regions (H), there were very few apoptotic cells. Unless stated otherwise, Blue: DAPI-labeled nuclei, Scale bar: 50 μm.

## Discussion

Biofabrication has emerged as one of the novel technologies for constructing tissues and organs in tissue engineering and regenerative medicine. The process holds promise for diseases and injuries that are currently difficult to treat due to lack of suitable therapies. In this study, we used a 3D-bioprinter to biofabricate structures from AMC-derived spheroids (Imamura et al., 2018; Moldovan et al., 2017). The structures were of a suitable size, shape, and autologous cellular composition to be transplanted into the cryo-injured urethras of rabbits. Thus, biofabricated structures have the potential to overcome limitations of the cell-injection methods. Additionally, we designed the biofabrications as C-shaped structures that would prevent urinary retention after transplantation. To our knowledge, the C-shaped structures are a unique and novel construct for a biofabricated structure and have not been reported.

AMCs that were attached and proliferating on the collagen-coated dishes were used to make the biofabricated structures. The cells were positive for the mesenchymal cell marker STRO-1, but they did not exhibit differentiation markers typical of muscle, nerve, and vascular cells that are present in the normal urethra. After the C-shaped structures were formed, the AMCs expressed integrin and/or cadherin, which are the cell-cell adhesion molecules. Cell-cell interactions play an important role in cell differentiation, and indeed, two and four weeks after transplantation, the biofabricated structures expressed markers typically found in normal urethras. Thus, the AMCs within the transplanted urethral biofabricated structures underwent differentiation into myoglobin-positive skeletal muscle cells, SMA-positive smooth muscle cells, S100- and CGRP-positive nerve cells, and vWF-positive vascular endothelial cells. These observations suggest that cell-cell interactions within the biofabricated structures might be a possible mechanism to enhance differentiation capacity after transplantation.

In addition to undergoing differentiation to form the highly specialized cells and tissues typical of the normal urethra, the AMCs in the transplanted tissue produced important signal molecules in the forms of growth factors and cytokines that are probably essential for cellular differentiation to occur. For instance, VEGF, which promotes myogenesis, muscle regeneration, and vascularization, (Frey et al., 2012; Levenberg et al., 2005) was present, as was NGF, which could promote reconstruction of nerve cells and/or formation of layered muscle structures (Lavasani et al., 2006). In addition, there might be direct or indirect roles for TGF-β1 and TNF-α that were present within the cells of the injured regions during the AMC-mediated reconstruction of the urethral muscle cells, blood vessels and nerve cells. The cytokine TGF-β is important in muscle regeneration in the absence of TNF-α, (Wynn and Vannella, 2016) while TNF-α alone promotes proliferation and fusion of myoblasts (Ji et al., 1998). However, in our study the transplanted cells contained both cytokines, suggesting the presence of a strong paracrine effect in which the two cytokines could act synergistically. Moreover, some cells that were originally surrounding the transplanted regions also produce growth factor and cytokines. Rapid conversion of the proinflammatory TNF-α-producing macrophages to an anti-inflammatory TGF-β-producing phenotype may be critical to the long-term survival of stem and progenitor cell populations in most tissues (Freytes et al., 2013; Heredia et al., 2013; Lemos et al., 2015). Furthermore, some AMCs from the transplanted biofabricated structure migrated from the transplanted regions into the tissues surrounding the transplanted areas and helped in reconstruction of the injured areas (data not shown).

Cells derived from AMCs also expressed desmin or pax7 at two and four weeks after transplantation. Desmin is a muscle-specific intermediate filament protein present in early myogenesis and is essential for the structural integrity and function of muscle (Paulin and Li, 2004). Pax7-positive cells might also play roles in regeneration and integration of muscle cells within the reconstructed muscle layers.

Urethras transplanted with the C-shaped AMC-biofabricated structures did not develop strictures that could impede the voiding of urine. Within the sham-operated control urethras, collagen fibers invaded the spaces of the damaged skeletal muscle and smooth muscle layers. There were also numerous P4HB- and HIF1α-positive cells, indicating a significant degree of hypoxia in the damaged region. In contrast, in the transplanted urethras, the collagen fibers were located between the recovering skeletal and smooth muscle layers, and there were few P4HB- and HIF1α-positive cells present. In addition, the structure-transplanted urethras had fewer apoptotic cells compared to the sham-operated urethras. We did not quantify the number of P4HB- and HIF1α-positive cells or, for that matter, the number of skeletal and smooth muscle cells or other cell types in the sham-operated or the AMC-transplanted urethras. However, the absence of structural recovery in the sham-operated urethras and the presence of structural recovery in the transplanted urethras was verified by histological and immunohistological investigations. Therefore, we hypothesize that tissue recovery mediated by the AMC-transplants inhibits the formation of urethral strictures.

We were unable to measure urethral physiological functions in this study. The most readily measured function is the leak point pressure (LPP) at which the bladder pressure forces urine to leak through the urethra. However, in the sham-operated controls, the presence of strictures narrowed the urethras, thus artificially raising the LPP, especially in comparison to the transplanted urethras that did not develop strictures. Thus, comparison of LPPs for the control and transplanted animals was not valid. Nevertheless, recovery of the cryo-injured urethras by transplantation of the biofabricated C-shaped AMC structures was strongly supported by the gross anatomy and microanatomy observations.

In conclusion, we used 3-D printing to manufacture novel biofabricated C-shaped structures composed of AMCs. The AMCs within the structures transplanted into the cryo-injured rabbit urethras survived and promoted recovery through differentiation of muscle, nerve, and vascular endothelial cells. In addition, these cells contained growth factors and/or cytokines that likely enhanced myogenesis, neurogenesis, and neoangiogenesis. The development of urethral strictures during the recovery process was inhibited by the biofabricated AMC-transplants. Therefore, transplantation of biofabricated structures derived from AMCs has the potential to provide novel and effective treatments for urine storage or voiding dysfunctions associated with damaged urethras.

## Materials and Methods

### Animals

Eight female New Zealand White rabbits (2.5-3.0 kg; Japan SLC, Inc., Shizuoka, Japan) at postnatal week 10 were used in this study. The rabbits were treated in accord with the Guide for the Care and Use of Laboratory Animals published by the US National Institutes of Health. The Animal Ethics Committee of Shinshu University School of Medicine approved the protocol used in this study.

### Preparation of AMCs and spheroids for biofabrication of C-shaped structures

Rabbits were anesthetized by intramuscular injection of midazolam (1.5 mg/kg; Astellas, Tokyo, Japan) and medetomidine hydrochloride (0.5 mg/kg; Zenoaq, Orion Pharma, Orion Corporation Espoo, Finland) and further anesthetized by inhalation of 3% sevoflurane (Abbot Japan Co., LTD., Tokyo, Japan). Previously, we established methods of isolation and harvesting autologous adipose-derived cells (Silwal Gautam et al., 2014). Briefly, intraperitoneal adipose tissue (approximately 1 g) was harvested from an abdominal incision. The adipose tissue was digested with 0.2% collagenase type I (Wako Pure Chemical Industries, Ltd., Osaka, Japan) for 90 minutes at 37° C with moderate rotary shaking. After shaking, the solution was centrifuged at 1,000 rpm for 4 minutes, and then the cells were suspended in culture medium composed of 4.5 g/L glucose-Dulbecco’s modified Eagle’s medium (Gibco, Thermo Fisher Scientific K.K., Kanagawa, Japan) supplemented with 15% regular fetal bovine serum (Biowest, Nuaille, France) and 0.1% penicillin-streptomycin (Gibco). After passing through a cell strainer (Life Science, Corning, NY, USA), primary cultures of the suspended cells were established on type I collagen-coated 10-cm culture dishes (AGC Techo Glass, Shizuoka, Japan) and incubated at 37°C in humid air with 5% CO_2_ for 5-7 days. The medium was completely changed every day to wash off unattached cells. After reaching confluence, the attached and proliferating cells were sub-cultured to increase the number of cells in a Collagen Type I-coated T 225 cm^2^ Flask (AGC Techno Glass).

To track the attached and proliferating cells in passages 2 or 3, a lipophilic red fluorescent dye that binds to cell membranes, PKH26 (Sigma Aldrich, St. Louis, MO, USA), was applied according to the manufacturer’s protocol. The labeled cells were re-suspended with Procul AD^®^ that was designed for human adipose mesenchymal stem cells (Rohto Pharmaceuticals, Osaka, Japan). To form cell spheroids, 100-μl volumes of 4×10^4^ cells/well were seeded in ultra-low attachment, U-shaped, 96-well plates (3-5 plates for each rabbit, Sumitomo Bakelite, Tokyo, Japan) and allowed to form cell aggregates at 37°C in 5% CO_2_ for 5 days.

### Biofabrication of C-shaped AMC structures

Biofabrication of the AMC structures was performed on a 3D-bioprinter (Regenova, Cyfuse Biomedical K.K., Tokyo, Japan) designed to create 3D-structures consisting only of cells (Imamura et al., 2018; Moldovan et al., 2017). Briefly, single spheroids, which were composed of the PKH26-labeled AMCs, were recovered from each well of the 96-well plates, and mounted on a custom designed, C-shaped 9×9 microneedle array. The formation of each biofabricated structure required 234 spheroids. The mounted spheroids, which were held in place by the microneedle array, were cultured with circulating Procul AD^®^ and allowed to form spheroid-spheroid contacts at 37°C in 5% CO_2_ for 7 days. On the seventh day of culture, the microneedle array was removed from the assembled biofabricated C-shaped structure (Fig. 1A).

### Production of cryo-injured urethral tissues and transplantation of the biofabricated C-shaped AMC structures

The rabbits from which the adipose tissue was harvested were anesthetized as above. A urethral catheter (Fr. 8, Terumo, Yokohama, Japan) was inserted, and then the urethra was exposed through a midline lower abdominal incision. The anterior wall of the urethra was sprayed with liquid nitrogen (Cryo Pro®, Cortex Technology, Hudsund, Denmark) for 20 s (Fig. 1B). The frozen region naturally thawed within 2-3 minutes by body temperature and/or room temperature. In this study, we define the frozen and thawed area as the cryo-injured regions. The cryo-injured regions were incised until the urethral catheter was visualized (Fig. 1C). After incision, the autologous biofabricated C-shaped AMC structure was placed around the urethral catheter (Fig. 1D). The wounded edges of the urethral tissue were sewed to the inserted C-shaped structure to secure them together and prevent the structure from falling freely into the urethral lumen or from falling out of the lumen. Finally, the abdominal incision was closed, and the urethral catheter removed. Four other rabbits served as sham-operated controls in which a urethral catheter was inserted, and the urethra was exposed and cryo-injured. The injured region was incised until the urethral catheter was visualized, and then the wounded edges were sewn together without the insertion of a C-shaped structure. At two and four weeks after surgery, the urethras were harvested, and the rabbits were sacrificed with the injection of an overdose of pentobarbital sodium solution (Kyoritsu Seiyaku Co., Tokyo, Japan).

### Histological and immunofluorescence evaluation

At two and four weeks after transplantation or sham operation, the harvested urethras were fixed in 4% paraformaldehyde for 12 hours. Then, the fixed samples were embedded in paraffin and cut in 5-μm thick serial sections. The sections were deparaffinized with xylene, rehydrated with ethanol, and rinsed three times with phosphate-buffered saline. Each section was stained with Masson trichrome, picrosirious red, or ApopTag^®^ ISOL fluorescence apoptosis detection kit (Merck Millipore, Merck KGaA, Darmstadt, Germany) according to the manufacturer’s protocol.

For immunohistochemistry, some of the deparaffinized sections were immersed in 10 mM sodium citrate (pH 6.0) and microwaved at 100°C for 10 min for antigen retrieval. The sections were blocked with 1.5% non-fat milk in phosphate-buffered saline for 1 h at 4°C.

To detect differentiated cells derived from the AMCs within the biofabricated structures, each section was incubated with one of the following antibodies for 12 h at 4°C to identify the following cellular markers: myoglobin (1:200, rabbit polyclonal; Spring Bioscience, Inc., Pleasanton, CA, USA), smooth muscle actin (SMA, 1:100 mouse monoclonal; Progen Biotechnik GmbH, Heidelberg, Germany), desmin (1:200 rabbit polyclonal; Progen Biotechnik GmbH), pax 7 (1:1000, rabbit; Lifespan Biosciences, Inc., Seattle, WA, USA), S100 (1:100, mouse monoclonal; Abcam, Cambridge, United Kingdom), calcitonin gene-related peptide (CRGP, 1:800, guinea pig polyclonal; Progen Biotechnik GmbH), or vascular endothelial cell marker von Willebrand factor (vWF, 1:100 rabbit polyclonal; Abcam).

To detect the presence of growth factors and cytokines in the AMCs of the biofabricated structures, the sections were incubated for 12 h at 4°C with antibodies for each of the following markers: vascular endothelial growth factor (VEGF, 1:100, rabbit polyclonal; Bioss, Woburn, MA, USA), nerve growth factor (NGF, 1:500, rabbit polyclonal; Abcam), transforming growth factor-β1 (TGF-β1, 1:100, rabbit polyclonal; Abcam), and tumor necrosis factor-α (TNF-α, 1:100, rabbit polyclonal; Abcam). Other sections were similarly incubated antibodies for P4HB (1:50: mouse monoclonal, Novus Biologicals, Inc.) or HIF1α (1; 50, rabbit polyclonal, Proteintech Group, Inc., Rosemont, IL, USA) for 12 h at 4°C.

After the primary antibody reactions, the sections were incubated with the respective secondary antibodies consisting of donkey anti-rabbit, -mouse, or -guinea pig, each conjugated with Alexa Fluor 488 (1:250; Molecular Probes, Eugene, OR, USA) for 1 h at 4°C. After rinsing, they were counterstained with 5 μg/ml 4’, 6-diamidino-phenylindole dihydrochloride (DAPI; Molecular Probes).

Biofabricated structures that were not used in a transplantation operation were immunohistochemically stained as described above with antibody for mesenchymal cell marker; STRO-1 (1:100, animal poly- or mono-, R&D System, Inc., Minneapolis, MN, USA) and the extracellular markers anti-integrin β1 (1:50, rat monoclonal; R&D Systems, Inc.) and N-cadherin (1:100, rabbit polyclonal; Bioss).

## Acknowledgements

This study was supported by a Grant-in-Aid for Scientific Research (C) from Japan Society for the Promotion of Science (16K11043).

## Data availability

None

